# Neural correlates of audio-visual integration of socially meaningful information in macaque monkeys

**DOI:** 10.1101/2021.05.02.442333

**Authors:** Mathilda Froesel, Maëva Gacoin, Simon Clavagnier, Marc Hauser, Quentin Goudard, Suliann Ben Hamed

## Abstract

Social interactions rely on the ability to interpret semantic and emotional information, often from multiple sensory modalities. In human and nonhuman primates, both the auditory and visual modalities are used to generate and interpret communicative signals. In individuals with autism, not only are there deficits in social communication, but in the integration of audio-visual information. At present, we know little about the neural mechanisms that subserve the interpretation of complex social events, including the audio-visual integration that is often required with accompanying communicative signals. Based on heart rate estimates and fMRI in two macaque monkeys (*Macaca mulatta*), we show that individuals systematically associate affiliative facial expressions or social scenes with corresponding affiliative vocalizations, aggressive facial expressions or social scenes with corresponding aggressive vocalizations and escape visual scenes with scream vocalizations. In contrast, vocalizations that are incompatible with the visual information are fully suppressed, suggesting top-down regulation over the processing of sensory input. The process of binding audio-visual semantic and contextual information relies on a core functional network involving the superior temporal sulcus (STS) and lateral sulcus (LS). Peak activations in both sulci co-localize with face or voice patches that have been previously described. While all of these regions of interest (ROIs) respond to both auditory and visual information, LS ROIs have a preference for auditory and audio-visual congruent stimuli while STS ROIs equally respond to auditory, visual and audio-visual congruent stimuli. To further specify the cortical network involved in the control of this semantic association, we performed a whole brain gPPI functional connectivity analysis on the LS and STS cumulated ROIs. This gPPI analysis highlights a functional network connected to the LS and STS, involving the anterior cingulate cortex (ACC), area 46 in the dorsolateral prefrontal cortex (DLPFC), the orbitofrontal cortex (OFC), the intraparietal sulcus (IPS), the insular cortex and subcortically, the amygdala and the hippocampus. Comparing human and macaque results, we propose that the integration of audio-visual information for congruent, meaningful social events involves homologous neural circuitry, specifically, an emotional network composed of the STS, LS, ACC, OFC, and limbic areas, including the amygdala, and an attentional network including the STS, LS, IPS and DLPFC. As such, these networks are critical to the amodal representation of social meaning, thereby providing an explanation for some of deficits observed in autism.

## Introduction

Brain structure and function have evolved in response to social relationships, both within and between groups, in all mammals. For example, across species, brain size and gyrification has been shown to increase with average social group size (Fox et al., 2017; Shultz & Dunbar, 2010; Van Essen & Dierker, 2007), as well as meta-cognitive abilities (Devaine et al., 2017). Within a given species, functional connectivity within the so-called social brain has been shown to be stronger in macaques living in larger social groups (Mars et al., 2012). In this context, successful social interactions require the proper interpretation of social signals (Ghazanfar & Hauser, 1999), whether visual (body postures, facial expressions, inter-individual interactions) or auditory (vocalization).

In humans, the core language system is amodal, in the sense that our phonology, semantics and syntax function in the same way whether the input is auditory (speech) or visual (sign). In monkeys and apes, vocalizations are often associated with specific facial expressions and body postures (Parr et al., 2005). This raises the question of whether and how auditory and visual information are integrated to interpret the meaning of a given situation, including emotional state and functional behavioral responses. For example, macaque monkeys *scream* as an indication of fear, triggered by potential danger from conspecifics or heterospecifics. In contrast, macaques *coo* during positive social interactions, involving approach, feeding and group movement. To what extent does hearing a scream generate a visual representation of the individual(s) involved in such an antagonistic situation, as opposed to a positive social situation, and does seeing an antagonistic situation set up an expectation that screams, but not coos, will be produced?

Face, voice, and social scene processing in monkeys have been individually explored, to some extent, from the behavioural (Gothard et al., 2004, 2009; Rendall et al., 1996; Sliwa et al., 2011) and the neuronal point of view (Aparicio et al., 2016; Arcaro et al., 2017; Cohen et al., 2007; Eifuku, 2014; Gil-da-Costa et al., 2004, 2006; Hesse & Tsao, 2020; Issa & DiCarlo, 2012; Joly, Pallier, et al., 2012; Joly, Ramus, et al., 2012; Moeller et al., 2008; Ortiz-Rios et al., 2015; Petkov et al., 2008; Pinsk et al., 2005, 2009; Poremba et al., 2003, 2004; Romanski et al., 2005; Russ et al., 2008; Schwiedrzik et al., 2015; Sliwa & Freiwald, 2017; Tsao et al., 2003). Audiovisual integration during naturalistic social stimuli has recently been shown in specific regions of the monkey face-patch system (Khandhadia et al., 2021), the voice-patch system (Ghazanfar, 2009; Ghazanfar et al., 2005; Perrodin et al., 2014, 2015), as well as in the prefrontal voice area (Romanski, 2012). However, beyond combining sensory information, social perception also involves integrating contextual, behavioural and emotional information (Freiwald, 2020; Ghazanfar & Santos, 2004). In this context, how macaque monkeys associate specific vocalizations with specific social visual scenes based on their respective meaning has scarcely been explored. Our goal is to help fill this gap.

This study used video-based heart rate monitoring and functional magnetic resonance in awake behaving monkeys to show that rhesus monkeys (*Macaca mulatta*) systematically associate the meaning of a vocalization with the meaning of a visual scene. Specifically, they associate affiliative facial expressions or social scenes with corresponding affiliative vocalizations, aggressive facial expressions or social scenes with corresponding aggressive vocalizations, and escape visual scenes with scream vocalizations. In contrast, vocalizations that are incompatible with the visual information are fully suppressed, indicating a top-down regulation over the processing of sensory input. Providing evidence of a homology with humans (Haxby et al., 2002; Haxby & Gobbini, 2011), we further show, using a functional connectivity analysis, that this audio-visual association involves two functionally coupled networks, one involved in the emotional processing of social stimuli, and one involved in their cognitive and attentional assessment.

## Results

In the following, we investigate whether and how macaques associate visual and auditory stimuli based on their semantic content, and we characterize the neuronal bases underlying this audio-visual integration. We obtained neural and autonomic data from two macaques using functional magnetic resonance brain imaging and video-based heart rate tracking. Each task combined visual stimuli of identical social content with either semantically congruent or incongruent monkey vocalizations. On each block of trials, the monkeys could be exposed to either visual stimuli only, auditory congruent stimuli only, auditory incongruent stimuli only, audio-visual congruent stimuli or audio-visual incongruent stimuli, in a blocked design (Figure 1a). Importantly, paired blocked conditions shared the same auditory stimuli, but opposite social visual content (Figure 1b). We report group fMRI and group heart-rate analyses. All reported statistics are based on non-parametric tests.

**Figure 1:**
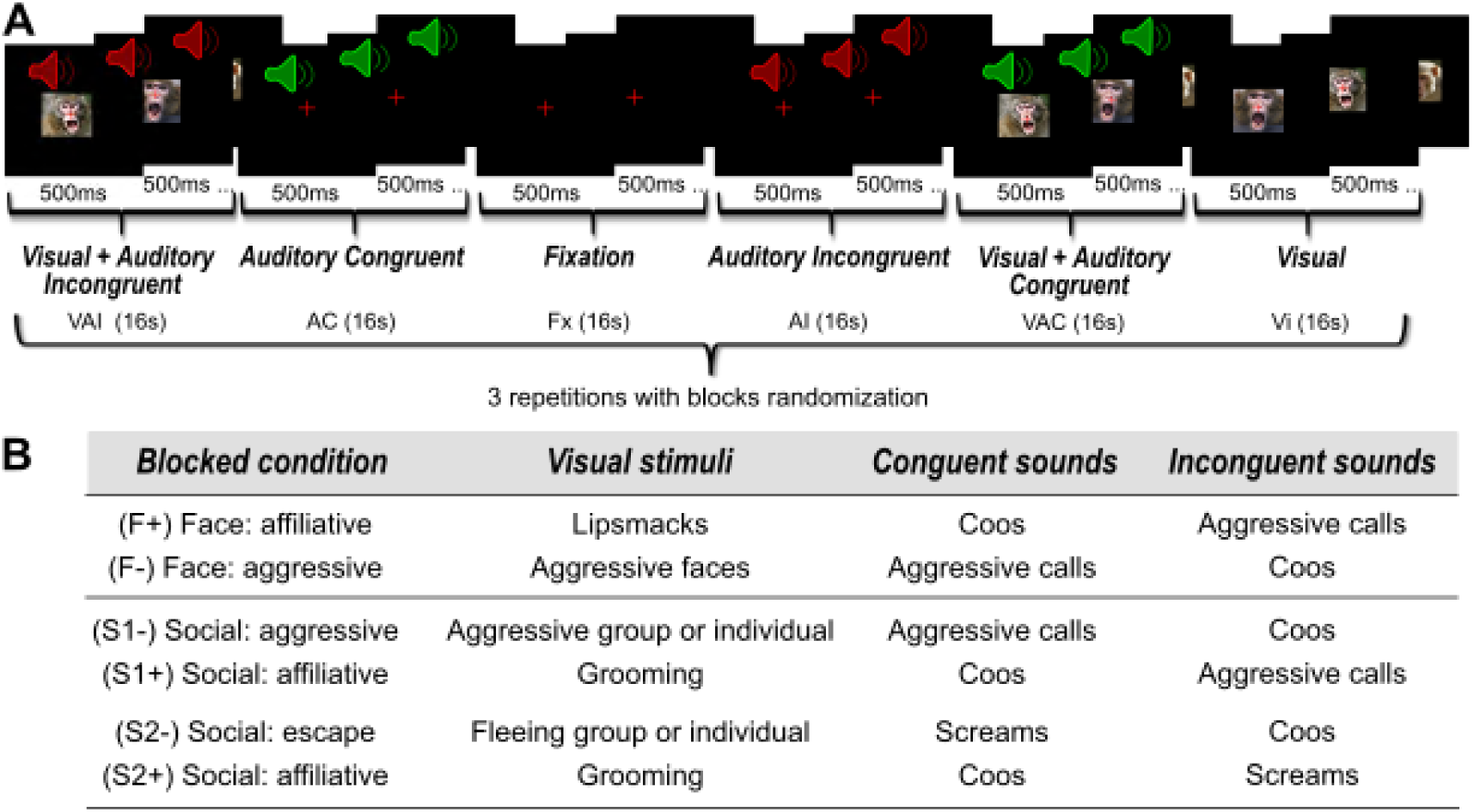
**A)** Experimental design. **Example** of an aggressive face (F-) blocked condition. Each run was composed of three randomized repetitions of six different blocks of 16 seconds. The six blocks could be visual (Vi), auditory with sounds congruent with the visual stimuli (AC), auditory with sounds incongruent with the visual stimuli (AI), audio-visual with sounds congruent with the visual stimuli (VAC), audio-visual with sounds incongruent with the visual stimuli (VAI), or fixation with no sensory stimulation (Fx). Each sensory stimulation block contained a rapid succession of 500ms stimuli. Each run started and ended with 10 seconds of fixation. **B) Description of blocked conditions**. Six different blocked conditions were used. Each blocked condition combined visual stimuli of identical social content with either semantically congruent or incongruent monkey vocalizations. Pairs blocked conditions shared the same auditory stimuli, but opposite social visual content (F+ vs. F-; S1+ vs. S1-; S2+ vs. S2-).

### Auditory whole brain activations depend on semantic congruence with visual context

Combining the F+ and F- face blocked conditions (Figure 2), which includes faces expressing lipsmacks or aggressive threats, we find robust bilateral activation (p<0.05 FWE) in the extra-striate cortex, along the superior temporal sulcus (STS) as well as in the prefrontal cortex, as expected from previous studies (Eifuku, 2014; Moeller et al., 2008; Tsao, Schweers, et al., 2008). Activations were also observed in the posterior part of the fundus of the intraparietal sulcus at an uncorrected level (p<0.0001). Please note that receiving coils were placed so as to optimize temporal and prefrontal cortex signal-to-noise ratio. As a result, no activations can be seen in the occipital cortex. The congruent auditory versus fixation contrast, which combined aggressive calls and coos in the two different blocked conditions, leads to activation within the inferior bank of the lateral sulcus, both at corrected (p<0.05 FWE) and uncorrected levels (p<0.0001), as described in previous studies (Joly, Pallier, et al., 2012; Petkov et al., 2008; Poremba et al., 2003). Importantly, this blocked condition also leads to the same robust bilateral activations as the visual contrast: the extra-striate cortex, along the superior temporal sulcus (STS) (p<0.05 FWE), as well as in the prefrontal and intraparietal cortex (p<0.0001 uncorrected). These activations are similar whether the congruent auditory stimuli are coos (Figure 3b) or aggressive calls (Figure 3c). In contrast, when we present the exact same aggressive calls and coos, the incongruent auditory versus fixation contrast leads to minimal activation, if any (Figure 2). Again, this doesn’t depend on whether the incongruent sounds are aggressive calls (Figure 3d) or coos (Figure 3e). This pattern of activation therefore confirms that auditory activation does not depend on the nature of the vocalization. Rather, it depends on whether the vocalizations are congruent or not to the semantic content of the visual stimuli.

**Figure 2:**
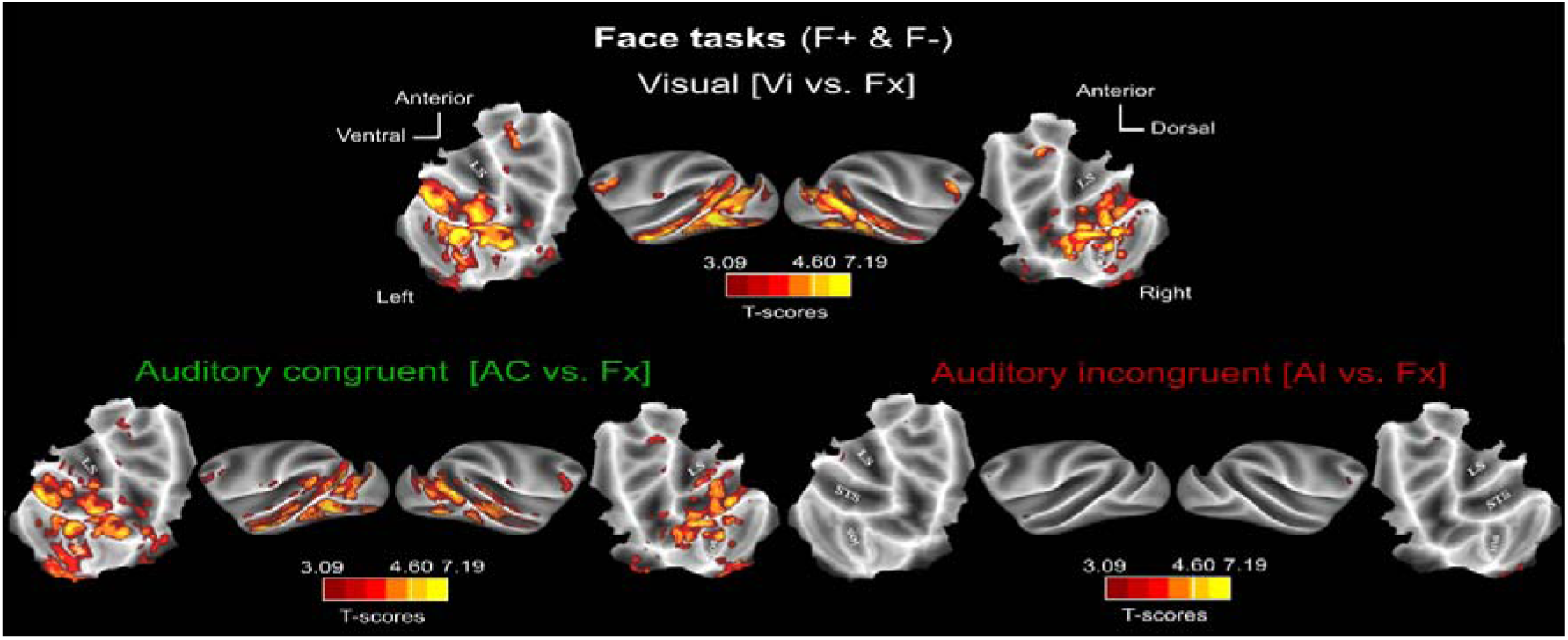
**Whole-brain activation FACE blocked condition (F+ & F-): main contrasts.** Whole-brain activation maps of the F+ (face affiliative) and F- (face aggressive) runs, cumulated over both monkeys, for the visual (white, Vi vs. Fx), auditory congruent (dark green, AC vs. Fx) and auditory incongruent (dark red, AI vs. Fx). Note that the AC and AI conditions contain exactly the same sound samples (coos and aggressive calls). Darker shades of red indicate level of significance at p<0.001 uncorrected, t-score 3.09. Lighter shades of yellow and brown outlines indicate level of significance at p<0.05 FWE, t-score 4.6. ios: Inferior Occipital Sulcus; LS: Lateral Sulcus; STS: Superior Temporal Sulcus.

**Figure 3:**
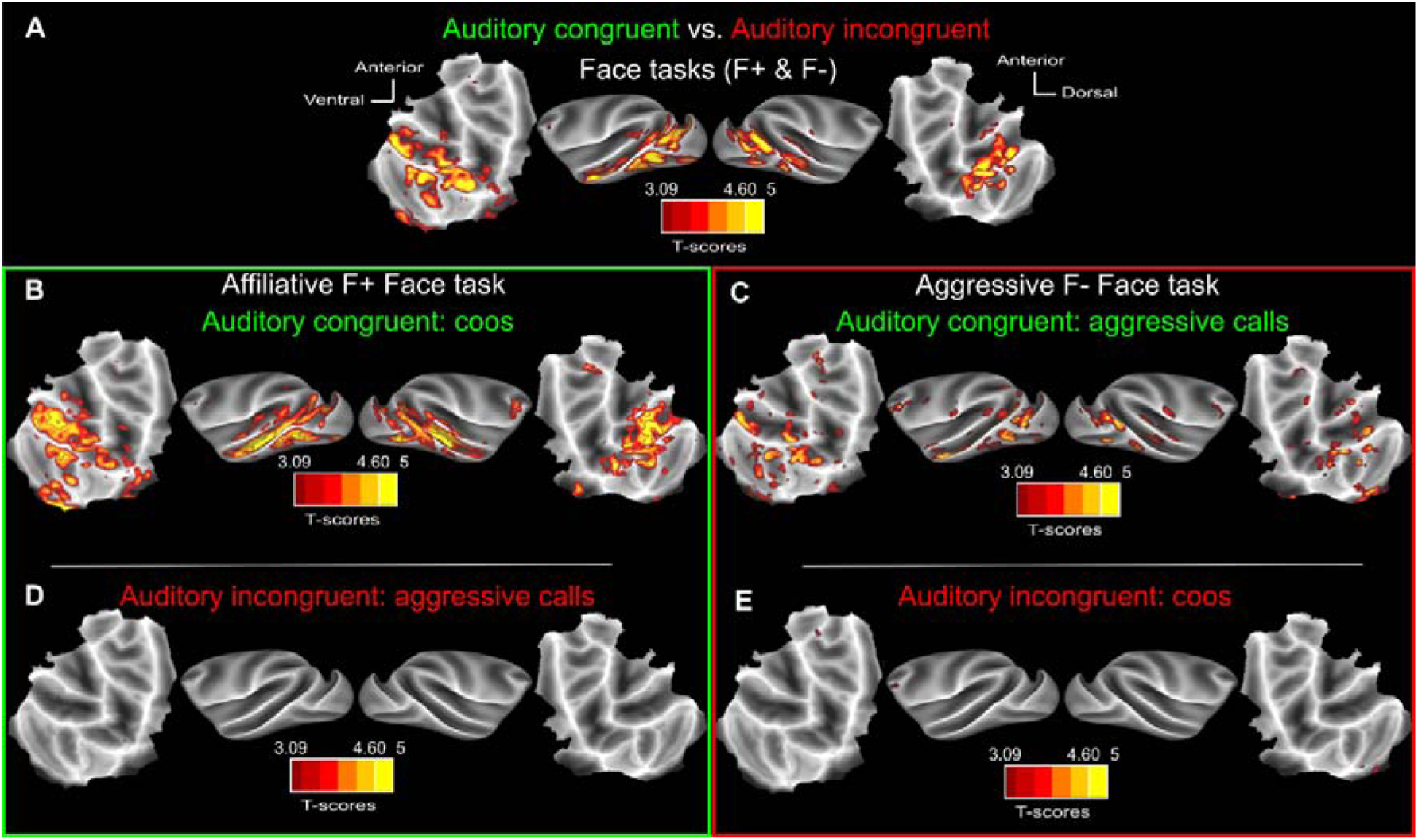
**Auditory activations depend on semantic congruence with visual context.** A) Whole-brain activation maps of the F+ (face affiliative) and F- (face aggressive) runs, for the *auditory congruent* vs *auditory incongruent* (relative to the visual context) contrast. Whole-brain activation map for the F+ (face affiliative) B) auditory congruent (coos, dark green, AC vs. Fx) and D) auditory incongruent (aggressive calls, dark red, AI vs. Fx) conditions. Whole-brain activation map for the F- (face aggressive) C) auditory congruent (aggressive calls, dark green, AC vs. Fx) and E) auditory incongruent (coos, dark red, AI vs. Fx) conditions. Darker shades of red indicate level of significance at p<0.001 uncorrected, t-score 3.09. Lighter shades of yellow and brown outlines indicate level of significance at p<0.05 FWE, t-score 4.6.

These observations are reproduced in a different set of blocked conditions, in which the visual stimuli involve social scenes (grooming, aggression or escape) with either semantically congruent or incongruent vocalizations (Figure 4 for all social blocked conditions and Figure S1 for S+ and S- social blocked conditions independently).

**Figure 4:**
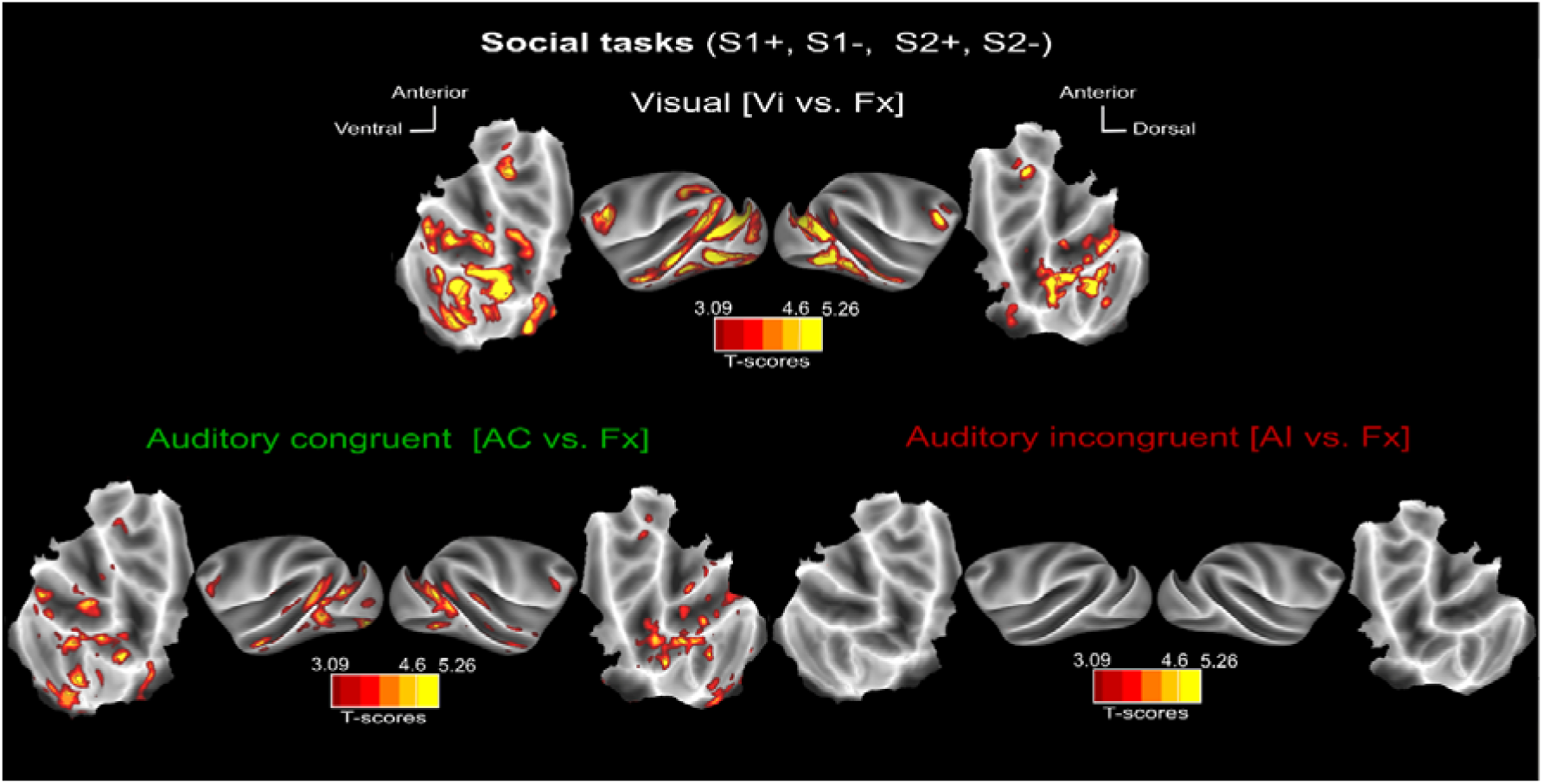
**Whole-brain activation Social blocked conditions (S1+, S1-, S2+ & S2-): main contrasts.** Whole-brain activation maps of the S1+, S2+ (social affiliative 1 & 2), S1- (social aggressive) and S2- (social escape) runs, cumulated over both monkeys, for the visual (white, Vi vs. Fx), auditory congruent (dark green, AC vs. Fx) and auditory incongruent (dark red, AI vs. Fx). Note that the AC and AI conditions contain exactly the same sound samples (coos, aggressive calls and screams). Darker shades of red indicate level of significance at p<0.001 uncorrected, t-score 3.09. Lighter shades of yellow and brown outlines indicate level of significance at p<0.05 FWE, t-score 4.6.

Taken together, these results indicate that audio-visual semantic associations are implemented in a specific cortical network involved in the processing of both visual face and social stimuli as well as auditory voice stimuli. An important question is thus whether these neuronal computations impact the behaviour or the physiology of the monkeys. In the following section, we investigate how heart rate changes in response to auditory-visual stimuli that are either congruent or incongruent with the social situation.

### Heart rate variations depend on semantic congruence with visual context

In this study, monkeys were required to fixate the centre of the screen while the different auditory and visual stimuli were presented. As a result, it was not possible to analyse whether gaze is spontaneously affected by the different stimulus categories. It was, however, possible to analyse heart-rate variation using a video-based method developed by our team (Froesel et al., 2020). Figure 5 focuses on heart rate variation in response to the auditory sound categories in the different blocked conditions. Although heart rate measures vary from one blocked condition to the other, in all blocked conditions, congruent auditory (Figure 5a, green) is systematically associated with lower heart rates than incongruent blocked condition (Figure 5a, red, Wilcoxon paired non-parametric test, p<0.001 for all blocked conditions except the F-task, p<0.05). This effect is more pronounced for the social blocked conditions (S1+/S1- and S2+/S2-) than for the face blocked conditions (Figure 5b, F+/F-, Friedman nonparametric test, p<0.001, N=127, Wilcoxon p<0.001), suggesting an intrinsic difference between the processing of faces and social scenes. This effect is also more pronounced for blocked conditions involving affiliative visual stimuli (F+, S1+ and S2+) than for blocked conditions involving aggressive or escape visual stimuli (Figure 5b, F-, S1- and S2-, Wilcoxon non-parametric test, p<0.001). This latter interaction possibly reflects an additive effect between the semantics and emotional valence of the stimuli. Indeed, affiliative auditory stimuli are reported to decrease heart rate relative to aggressive or alarm stimuli (Kreibig, 2010). As a result, emotionally positive stimuli would enhance the semantic congruence effect, while emotionally negative stimuli would suppress the semantic congruence effect. Overall, these observations indicate that semantic congruence is perceptually salient, at least implicitly.

**Figure 5:**
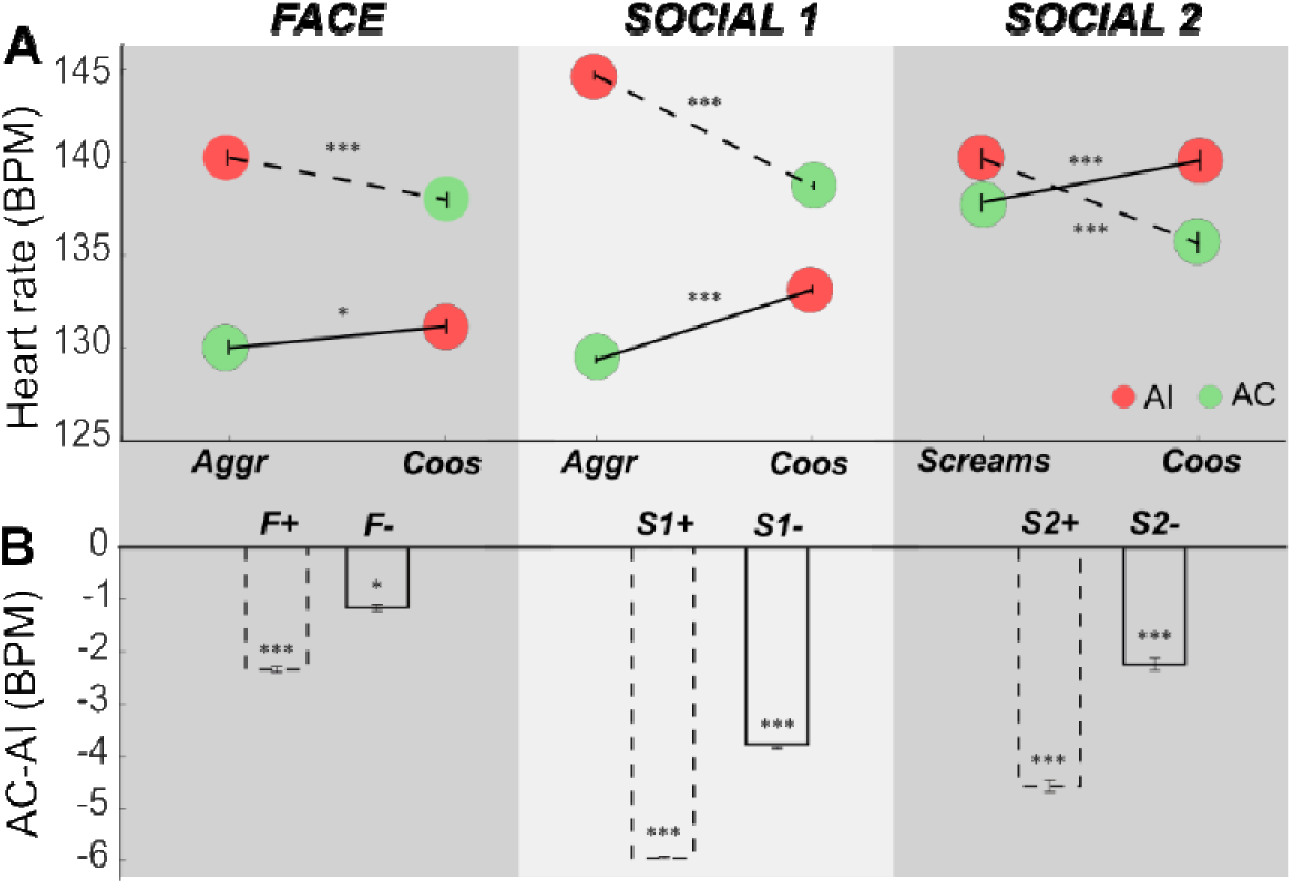
**Blocked condition-related heart rate (BMP) variations.** A) Absolute heart rate (BMP, beats per minute) during the congruent (green) and incongruent (red) auditory blocks of each task. Dashed lines correspond to the affiliative blocked condition as defined by the visual stimuli, whereas continuous lines refer to aggressive or escape blocked conditions. Blocked conditions are defined by pairs involving the same vocalization categories but different visual stimuli, as defined in Figure 1b. Each blocked condition pair shows significantly higher heart rates for incongruent auditory stimuli compared to congruent auditory stimuli (Friedman nonparametric test, Face: p<0.001, 271.442, N=254; Social 1: p<0.001, 295.34, N=254; Social 2: p<0.001, 174.66, N=254). This is also true for each individual blocked condition (Wilcoxon paired non-parametric test, p<0.001 for all blocked conditions except F-: p<0.05), B) Difference between AC and AI bloc means. All significantly different from zero (Wilcoxon paired non-parametric test, p<0.001 for all blocked conditions except F-: p<0.05).

### Visual auditory gradients across the lateral sulcus (LS) and superior temporal sulcus (STS)

While LS demonstrates stronger activation for socially congruent auditory stimuli relative to visual stimuli, the STS appears to be equally activated by both sensory modalities. To better quantify this effect, we define regions of interest (ROIs, 1.5 mm spheres) at local peak activations in the auditory congruent (AC vs Fx) contrast, in the face (F+ and F-) blocked conditions (Figure 6A, see Figure S2 for a precise localization of each of these local maxima on corresponding brain anatomy). These peaks match peak activations in the social blocked conditions (S1+, S1-, S2+ and S2-) auditory congruent (AC vs Fx) contrast. This latter social blocked condition contrast reveals two additional peaks in the right LS which were used to define two additional ROIs (right LS4 and LS6). Overall, 8 ROIs are thus defined in the right STS, 6 in the left STS, 4 in the left LS and 6 in the right LS. The numbering of these ROIs was adjusted so as to match mirror positions across hemispheres. Figure 6B presents mean percentage signal change (%PSC) for each independent ROI, in the left and right hemispheres, on each of the 5 face and social blocked conditions respectively. Overall, STS ROIs and LS ROIS had similar %PSC profiles across the 5 blocked conditions for each group of blocked conditions (face vs. social). No interhemispheric difference could be noted.

**Figure 6:**
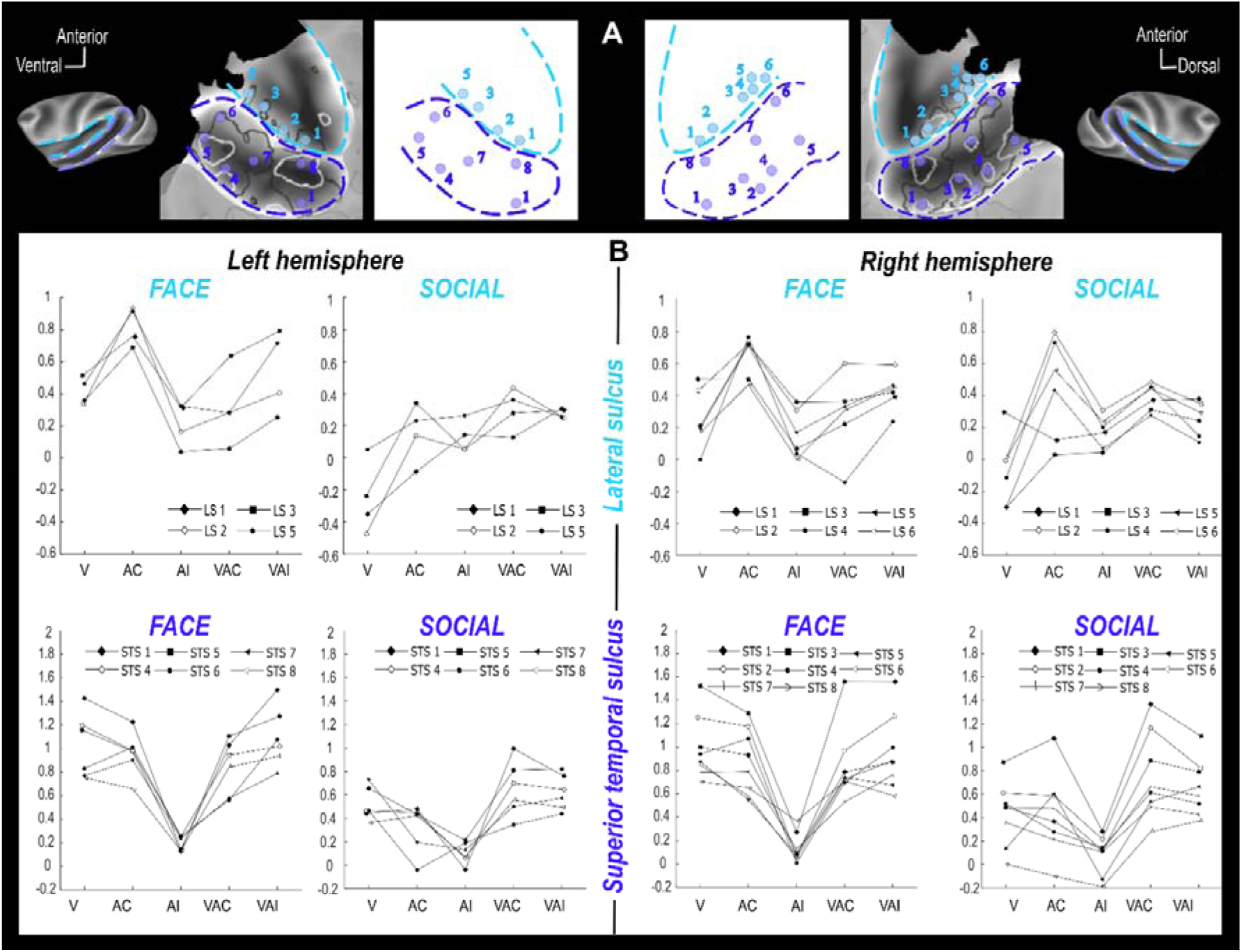
**Percentage of signal change (%PSC) for selected left and right hemisphere ROIs in the lateral sulcus (light blue) and in the superior temporal sulci (dark blue). (A)** ROIs are 1.5mm spheres located at local peak activations. Left and right hemisphere numbering associate mirror ROIs. ROI location in the each of the left and right STS and LS is described in the bottom flat maps. (B) %PSC (mean) are presented for each ROI (8 in right STS, 6 in left STS, 4 in left and 6 in right lateral sulcus) and each blocked condition of interest (V: visual, AC: auditory congruent, AI: auditory incongruent, VAC: visuo-auditory congruent, VAI: visuo-auditory incongruent).

In the STS, in both of the face (F+ and F-) and social blocked conditions (S1+, S1-, S2+ and S2-), %PSC in the visual blocked condition relative to fixation across all ROIs is not significantly different from %PSC in the auditory congruent blocked condition relative to fixation, (Figure 7, left, Wilcoxon non-parametric test). The STS thus appears as equally responsive to visual and auditory social stimuli (%PSC of all blocked conditions are significantly different from fixation %PSC, Wilcoxon non-parametric test, p<0.01 or p<0.001). In contrast, in the LS, %PSC in the visual blocked condition relative to fixation across all ROIs is significantly different from %PSC in the auditory congruent blocked condition relative to fixation, (Figure 7, left, Wilcoxon non-parametric test, p<0.005). This result therefore suggests a strong auditory preference for LS (%PSC of all auditory are significantly different from fixation %PSC, Wilcoxon non-parametric test, p<0.01), although LS is also significantly activated by the visual stimuli in the face blocked condition (p<0.01). Overall, therefore, LS appears preferentially sensitive to auditory stimuli while STS appears to be equally responsive to visual and auditory stimuli.

**Figure 7:**
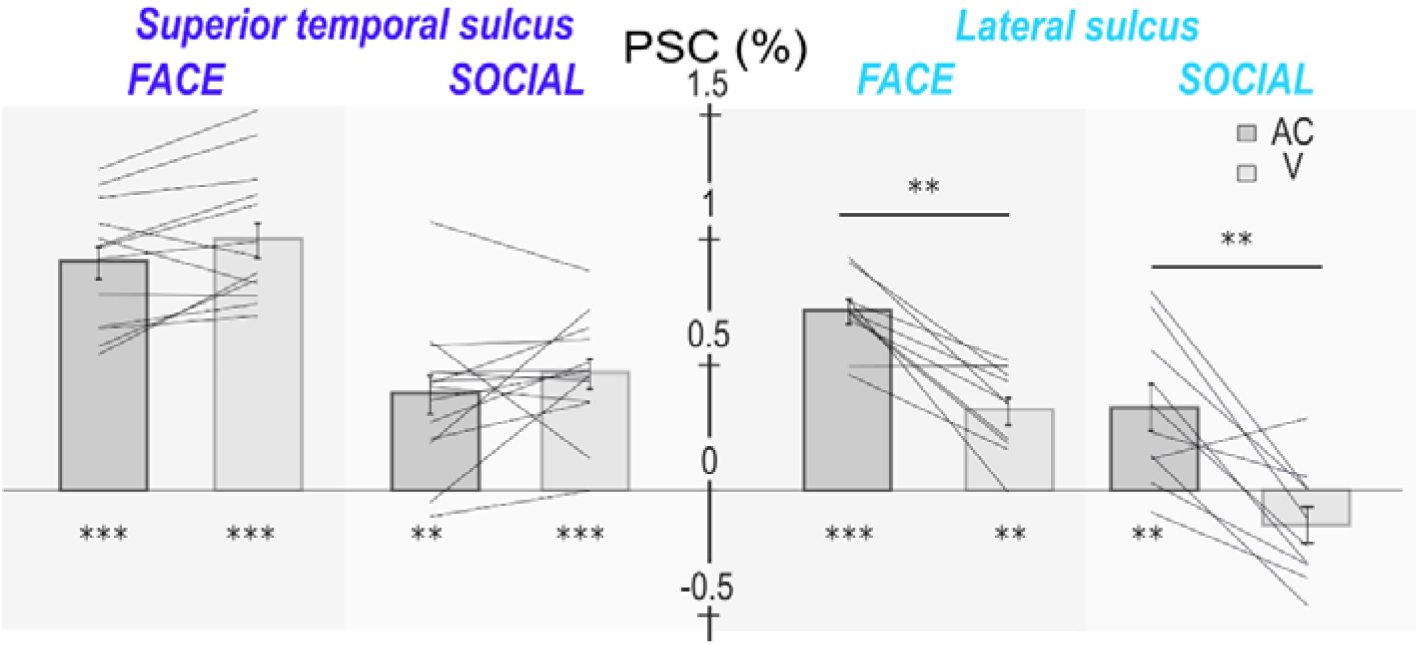
**Percentage of signal change (%PSC) across all lateral sulcus (light blue) and superior temporal sulci (dark blue) ROIs of both hemispheres, comparing the auditory and visual blocked conditions.** Statistical differences relative to fixation are between blocked conditions and indicated as follows: ***, p<0.001; **, p<0.01 (Wilcoxon non-parametric test).

### Visual-auditory integration in the STS during the social blocked conditions

When processed in the brain, sensory stimuli from different modalities are combined such that the neuronal response to their combined processing is different from the sum of the neuronal responses to each one of them. This process is called multisensory integration (Avillac et al., 2007) and is more pronounced when unimodal stimuli are ambiguous or difficult to perceive (Alais & Burr, 2004; Ernst & Banks, 2002). The question here, therefore, is whether and how the LS and the STS combine visual and auditory social stimuli as a function of their semantic congruence. Multisensory integration is not straightforward to assess based on fMRI signals. A minimal criterion here would be to have significant %PSC signal differences between the bimodal blocked conditions and both of the unimodal blocked conditions. Figure 8 shows the whole brain activation maps obtained for the visual-auditory blocked condition contrasted, with fixation, the visual blocked condition and the auditory blocked condition, for the congruent (Figure 8A) and incongruent (Figure 8B) auditory vocalizations, for the face (Figure 8, left panel) and the social blocked conditions (Figure 8, right panel). Figure 8C presents the contrast between the congruent and incongruent visuo-auditory blocked conditions.

**Figure 8:**
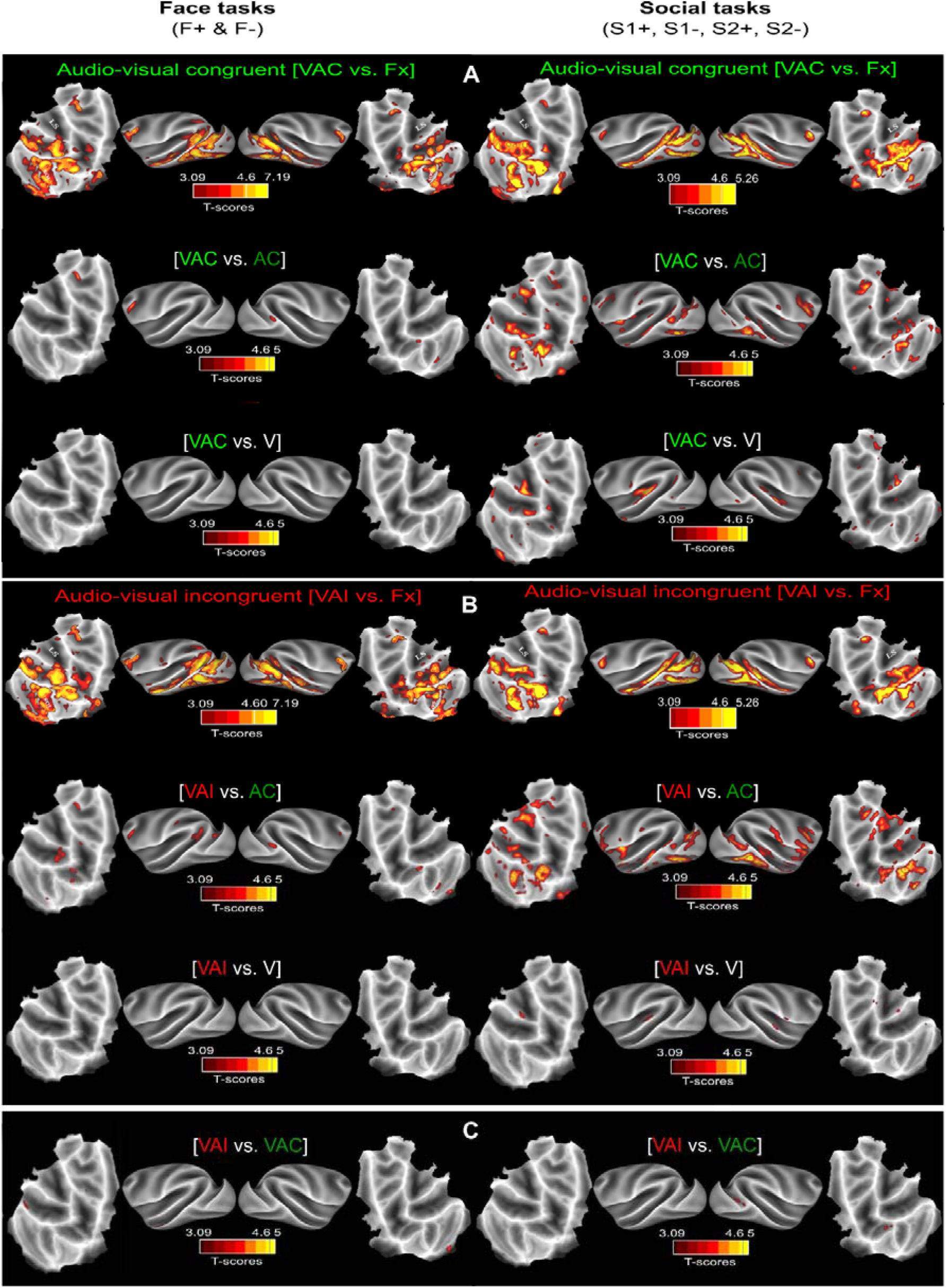
**Whole-brain activations for the Face (F+& F-) and Social blocked conditions (S1+, S1-, S2+ & S2-): bimodal versus unimodal contrasts.** A) Whole-brain activation maps of the F+ (face affiliative) and F- (face aggressive) runs (left panel) and the S1+, S2+ (social affiliative 1 & 2), S1- (social aggressive) and S2- (social escape) runs (right panel), for the congruent auditory vocalizations (green). Contrasts from top to bottom: audio-visual vs. fixation, audio-visual vs. auditory and audio- visual vs. visual. B) Same as in A) but for the incongruent auditory vocalizations (red). C) Whole-brain activation maps for the audio-visual incongruent vs audio-visual congruent contrast. All else as in A). Darker shades of red indicate level of significance at p<0.001 uncorrected, t-score 3.09. Lighter shades of yellow and brown outlines indicate level of significance at p<0.05 FWE, t-score 4.6.

Overall, in the face blocked condition, activations in the audio-visual conditions are not significantly different from the visual and auditory conditions alone (Figure 8A&B, left panel). Likewise, no significant difference can be seen between the congruent and incongruent visuo-auditory conditions (Figure 8C, left panel). Figure S3 compares %PSC for the bimodal and unimodal conditions across all STS selected ROIs and all LS selected ROIs. Neither reach the minimal criteria set for multisensory integration. In the social blocked condition, activation in the audio-visual conditions show local significant differences relative to the visual and auditory conditions alone (Figure 8A&B, left panel), but none in the same regions. However, when comparing the %PSC for the bimodal and unimodal conditions across all STS selected ROIs and all LS selected ROIs, the STS ROIs reach the minimal criteria set for multisensory integration, as their %PSC is significantly different from each of the bimodal conditions and each of the unimodal conditions (Wilcoxon non-parametric test, p<0.01). Thus multisensory integration appears to take place, specifically in the STS, and during the social blocked conditions, possibly due to the higher ambiguity in interpreting social static scenes relative to faces (Figure S4). Importantly, and while most significant activations in the bimodal vs. unimodal auditory condition are located within the audio-visual vs. fixation network, a bilateral activation located in the anterior medial part of the LS deserves attention. Indeed, this activation, encompassing part of the insula and of anterior SII/PV, is identified both in the congruent and incongruent auditory conditions and might be involved in the interpretation of semantic congruence between the visual and auditory stimuli. This possibility is addressed next.

### Semantic visuo-auditory association network

Both our whole brain fMRI analyses and heart rate measures indicate that the monkeys actively associate the auditory voice stimuli with the social context set by the visual stimuli, whether faces or social scenes. T to further specify the cortical network involved in the control of this semantic association, we performed a whole brain gPPI functional connectivity analysis on the LS and STS cumulated ROIs, to identify the cortical and subcortical regions that systematically correlate with either the LS or STS time series, irrespective of the blocked condition. To increase the statistical power of our analysis, we independently performed this gPPI analysis on each individual task. Table 1 of the Supplementary Material summarizes for selected cortical regions, the blocked conditions and ROIs for which a significant correlation is observed in at least one voxel (p<0.005 uncorrected). In the following, we only consider for further discussion the cortical regions that correlate, with either the LS (Figure 8a) or STS time series (Figure 8b) in: (criteria-1) at least three of the six blocked conditions, for each of the left and right hemispheres; (criteria-2) at least three of the six blocked conditions in at least one hemisphere and in at least eight of the blocked conditions across both hemispheres (Figure 8, bold front). This gPPI analysis highlights a functional network connected to the LS and involving the anterior cingulate cortex (ACC), area 46 in the dorsolateral prefrontal cortex (DLPFC), the orbitofrontal cortex (OFC), the intraparietal sulcus (IPS), the superior temporal sulcus (STS), and subcortically, the amygdala and the hippocampus. The same core network is identified for its connectivity with the STS, with the addition of the insula. It is worth noting that the dorsal pulvinar, although at the limit of the inclusion criteria, was detected several times. This nucleus often acts as a attentional modulator and is implicated not only in social perception, but can also be considered as an audio-visual integrator given its connection with face patches (Schwiedrzik et al., 2015), and the auditory and visual cortex (Froesel et al., 2021). Further investigation should be carried out to determine its role in this network.

## Discussion

Based on heart rate estimates and fMRI, our results show that rhesus monkeys systematically associate affiliative facial expressions or social scenes with corresponding affiliative vocalizations, aggressive facial expressions or social scenes with corresponding aggressive vocalizations, and escape visual scenes with scream vocalizations. In contrast, vocalizations that are incompatible with the visual information are fully suppressed, suggesting a top-down regulation over the processing of sensory input. In other words, rhesus monkeys correctly associate the meaning of a vocalization with the meaning of a visual scene. This audio-visual, semantic binding with contextual information relies on a core functional network involving the superior temporal sulcus (STS) and the lateral sulcus (LS). LS regions of interest (ROIs) have a preference for auditory and audio-visual congruent stimuli while STS ROIs respond equally to auditory, visual and audio-visual congruent stimuli. A functional connectivity analysis (gPPI) identified a functional network connected to the LS and STS, involving the anterior cingulate cortex (ACC), area 46 in the dorsolateral prefrontal cortex (DLPFC), the orbitofrontal cortex (OFC), the intraparietal sulcus (IPS), the insula and subcortically, the amygdala and the hippocampus. Overall, we propose that the integration of congruent social meaning from audio-visual information involves an emotional network composed of the STS, LS, ACC, OFC, and limbic areas, including the amygdala, and an attentional network including the STS, LS, IPS and DLPFC. These observations are highly robust as they are reproduced over six sets of independent behavioral blocked conditions, involving distinct associations of visual and auditory social information.

### Interpretation of social scenes and vocalization by macaque monkeys

As is the case for human oral communication, monkey vocalizations are expected to be interpreted as a function of their emotional or contextual meaning. For example, a monkey scream indicates potential danger, is associated with fear and calls for escape and flight from the dangerous context. In contrast, coos are produced during positive social interactions and often elicit approach. Here, we show that when two different types of vocalizations are presented together with a social visual stimulus, the heart rate of the monkeys significantly decreases when the vocalization is congruent with the visual scene as opposed to incongruent. Likewise, we show that the activity of the voice processing network is dramatically suppressed in response to the incongruent vocalization. This pattern of activation provides direct neurobiological evidence that macaques infer meaning from both social auditory and visual information and are able to associate congruent information. In the network of interest, activations are not significantly different between the auditory, visual or audio-visual conditions. Most interestingly, aggressive calls are associated with both aggressive faces and aggressive social scenes, whereas coos are associated with both lipsmacks and inter-individual social grooming. We thus propose that these networks might represent social meaning irrespective of sensory modality, thereby implying that social meaning is amodally represented. We hypothesize that such representations are ideal candidate precursors to the lexical categories that trigger, when activated, a coherent set of motor, emotional and social repertoires.

### Audio-visual social stimuli robustly activate the face and voice patches

Face processing is highly specialized in the primate brain (Hesse & Tsao, 2020). In the macaque brain, it recruits a specific system called the face patch system, composed of interconnected areas, identified by both fMRI (Afraz et al., 2015; Aparicio et al., 2016; Arcaro et al., 2017; Eifuku, 2014; Freiwald & Tsao, 2010; Hadj-Bouziane et al., 2008; Issa & DiCarlo, 2012; Moeller et al., 2008; Pinsk et al., 2005, 2009; Tsao et al., 2003) and single cell recording (Grimaldi et al., 2016; Moeller et al., 2008; Tsao et al., 2006). This system recruits areas in the superior temporal sulcus, as well as in the prefrontal and orbito-frontal cortex. Specific limbic and parietal regions are also recruited together with this core system during, respectively, the emotional and attentional processing of faces (Schwiedrzik et al., 2015). The core face patches are divided into five STS areas (Anterior medial, AM; anterior fundus, AF; anterior lateral, AL; middle fundus, MF and middle lateral ML) and the PL (posterior lateral patch), a posterior face patch in the occipital cortex (Eifuku, 2014; Hesse & Tsao, 2020; Tsao et al., 2003; Tsao, Moeller, et al., 2008). Based on a review of the literature, and anatomical landmark definitions, we associate the activation peaks identified in the present study with these five face patches (Figure 10). Correspondence is unambiguous and the STS 4 ROIs matches ML, STS 7 matches MF, STS 5 matches AL and STS 6 matches AF. The occipital face patch PL is also identified in the general contrast maps as well as the frontal area defined in the literature as PA (prefrontal accurate) (Tsao, Schweers, et al., 2008). It is worth noting that in our experimental design, these face patches are activated both during the purely auditory congruent condition as well as during the visual conditions. Such activations are not reported during purely auditory conditions, indicating that this network is recruited during audio-visual association based on meaning.

**Figure 9:**
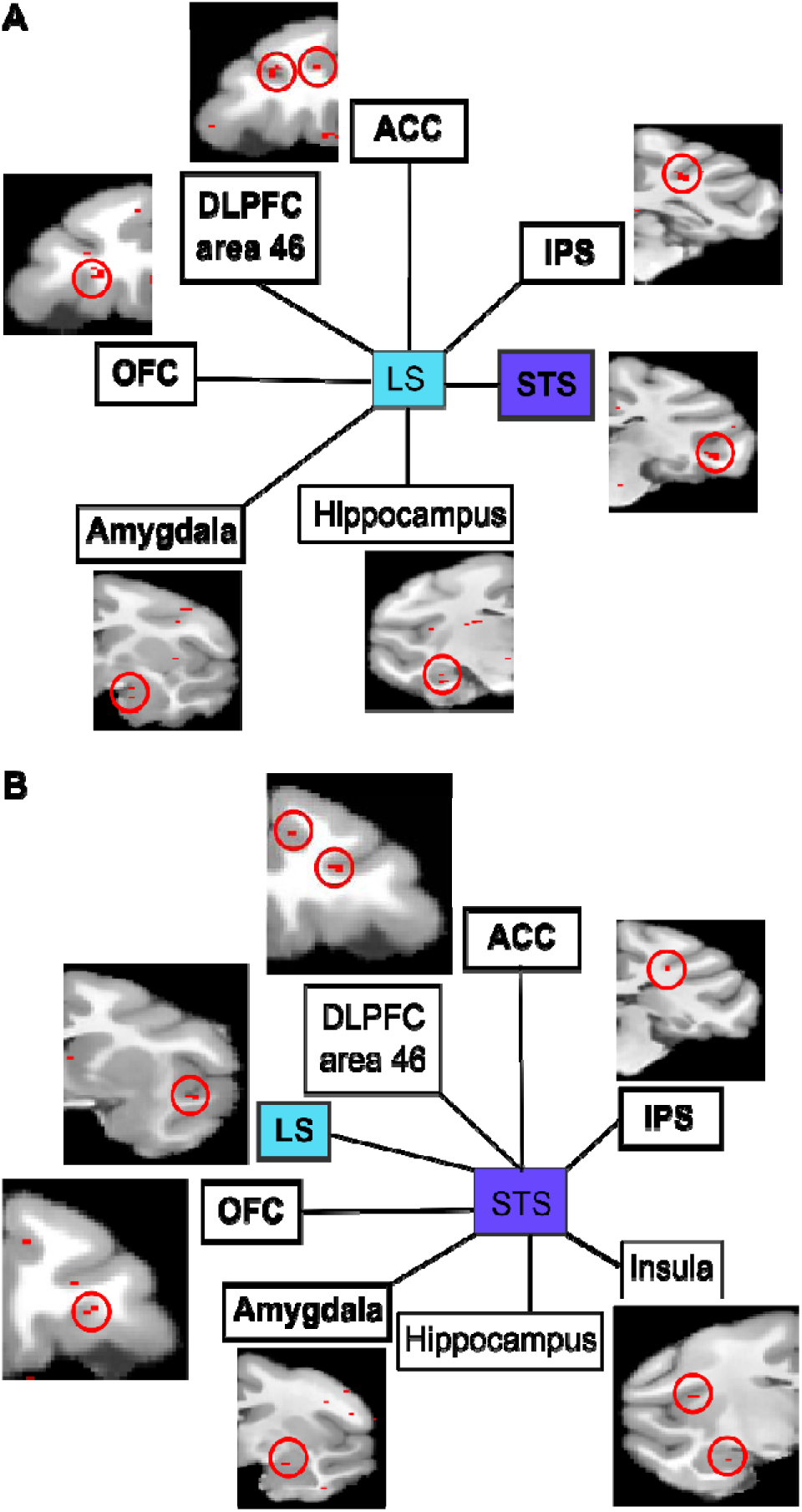
**Whole brain gPPI functional connectivity analysis for the lateral sulcus (A) and in the superior temporal sulcus ROIs (B).** Reported here are functional connectivity analyses between cortical regions under the following statistical criteria. Areas in bold fonts show a significant correlation at p<0.005 uncorrected in at least 4 out of the 6 individual blocked conditions, in each hemisphere (see supplementary table 1). Areas in regular font show a significant correlation at p<0.005 uncorrected in at least 3 out of the 6 individual blocked conditions, in each hemisphere (see supplementary table 1). ACC: Anterior Cingulate Cortex; DLPFC: dorsolateral prefrontal cortex; OFC: orbitofrontal cortex; IPS: Intraparietal sulcus.

**Figure 10:**
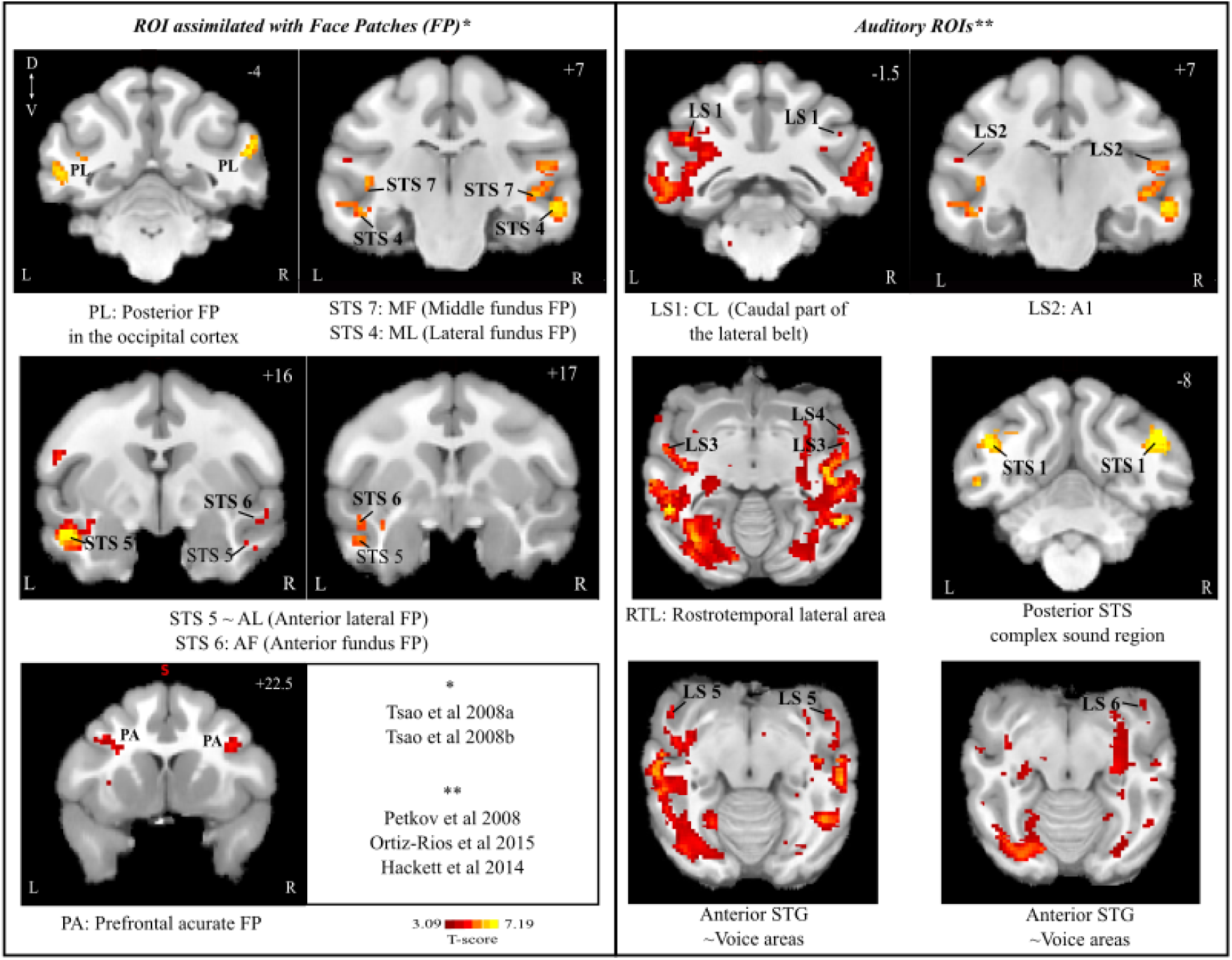
**Correspondence between task-related ROIs and face patches (left panels) and voice areas (right panels).** Color-scale runs start at p<0.001 uncorrected levels. Task related ROIs are numbered as in Figure 5. PA (prefrontal acurate); AM (anterior medial); AF (anterior fundus); AL (anterior lateral); MF (middle fundus); ML (middle lateral); PL a posterior face patch in the occipital cortex; CL (Caudal part of the lateral belt), A1 (primary auditory cortex), RTL (Rostrotemporal lateral area), STS (Superior Temporal Sulcus), STG (Superior Temporal Gyrus). *: Sources for face patch localization. **: Sources for voice areas.

Similarly to face patches, voice processing also involves a system composed of voice patches (for review, see Belin, 2017). In macaques, voice specific areas include the anterior superior temporal gyrus (aSTG), the orbitofrontal cortex (OFC) and a part of the STS close to the lateral sulcus (Cohen et al., 2007; Joly, Pallier, et al., 2012; Joly, Ramus, et al., 2012; Perrodin et al., 2015; Petkov et al., 2008; Poremba et al., 2003). The auditory processing circuit is proposed to be organized in two main networks, a ventral and a dorsal network (see for review Kuśmierek & Rauschecker, 2014), such that the auditory ventral stream is activated by species-specific vocalizations whereas the dorsal stream is involved in the spatial location of sounds (Ortiz-Rios et al., 2015; Russ et al., 2008). This functional dissociation is observable as early as in the lateral belt such that its caudal part (CL) is selectively associated with sound location while the anterior part (AL) is more linked to sound identity such as vocalizations (Kuśmierek & Rauschecker, 2009, 2014; Tian et al., 2001). Again, based on a review of the literature, and anatomical landmark definitions, we associate the activation peaks identified in the present study with these voice patches (Figure 10). Correspondence is unambiguous and the LS 1 ROI can be associated to CL (i.e. dorsal sound processing pathway) and LS2 to core primary auditory area A1. Within the ventral sound processing pathway, LS 4 ROI can be associated to area AL, LS 5 to rostro-temporal lateral area (RTL) and LS 6 to the rostro-temporal polar field (RTp). Last, LS 6 is compatible with the anterior most voice STG area described by Petkov and colleagues (2008). The voice patch system also involves the ventral dorsolateral prefrontal cortex or vlPFC (Romanski et al., 2005), located in the inferior dimple at the boundary between area 45a and 46 (Petrides & Pandya, 2002). This cortical region has been proposed to play a key role in the cognitive control of vocalizations as well as in the interpretation of call meaning (Romanski & Averbeck, 2009). Microstimulations further indicate that this prefrontal voice patch is functionally connected with the putative macaque homologue of human’s Broca area 44 (Rocchi et al., 2021). In the present study, the ventral prefrontal activation, while matching nicely with the PA face patch, only partially overlaps with the prefrontal voice patch, suggesting a possible functional specialization. Taken together, these results indicate that the association between vocalization meaning and social visual stimuli recruits the face and voice patch system.

In the right hemisphere, two supplementary STS activations are reported, STS 2 and STS 3. They are located posteriorly to the putative ML face patch and possibly coincide with the gaze following patch reported in the dorsal posterior infero-temporal cortex (PITd) (Marciniak et al., 2014). This cortical region is recruited during motion discrimination blocked conditions (Stemmann & Freiwald, 2016) and is proposed to play a key role in attentional selection due to its strong connectivity with the dorsal attentional areas such as lateral intraparietal areas (LIP) and frontal eye fields (FEF) (Sani et al., 2019). In spite of the fact that the monkeys did not have to produce any active behaviour, these activations possibly reflect enhanced, automatic, attention to social cues.

Visual fMRI activations have already been described in the LS, in the primary auditory cortex and in the non-primary core (belt) (Kayser et al., 2007). This observation has been confirmed using single cell recording studies (Kayser et al., 2008). In contrast, to our knowledge, no extended auditory responses have yet been described in the STS. This suggests that the STS auditory activations described here arise from the task design and the implicit association between a visual stimulus and two competing auditory stimuli. An important question to be addressed by single unit recording studies is whether these STS auditory activations correspond to neuromodulatory LFP modulations or to actual spiking activity. Quite interestingly, while we identify an audio-visual gradient between the LS and the STS, the LS showing higher activations for voice as compared to visual social stimuli, and the STS showing equal responses to both, no clear gradient of auditory or visual activations can be identified either within the STS or within the LS. This suggests that voice-social visual associations rely on the activity of the entire network, rather than on some of its subparts.

### Audio-visual association based on meaning and multisensory integration

The strict definition of multisensory integration involves the combination of sensory inputs from different modalities under the assumption of a common source (Lee & Noppeney, 2014; Stein et al., 2014). In this context, it has been shown that multisensory integration speeds up reaction times and enhances perception (Grant & Seitz, 2000; Lehmann & Murray, 2005; Murray et al., 2005; Raab, 1962; Welch et al., 1986), including when processing lip movement during speech (Navarra & Soto-Faraco, 2007; Shahin & Miller, 2009; Van Wassenhove et al., 2005). Multisensory processes are also at play to predict the consequences of one modality onto another, i.e. in the temporal domain (Cléry et al., 2015, 2017; Cléry et al., 2020; Guipponi et al., 2015). At the neuronal level, multisensory integration is defined as a process whereby the neuronal response to two sensory inputs is different from the sum of the neuronal responses to each on its own (Avillac et al., 2007; Stein et al., 2009). In the present study, the auditory and visual stimuli are associated based on their meaning (e.g., coos are associated with grooming) and possible contingency (e.g., screams are associated with escape scenes). In addition, incongruent auditory stimuli are actively suppressed by visual context. As a result, this association based on stimulus meaning does not correspond to the low level association classically understood by multisensory integration. Yet, one could expect that associating meaning might lead to an enhancement of neuronal processes similar to that described during multisensory integration. To probe this hypothesis, we apply the less stringent multisensory integration criteria used in fMRI studies, namely we test for audio-visual responses statistically higher (or lower) than each of the uni-sensory conditions (Beauchamp, 2005; Gentile et al., 2010; Pollick et al., 2011; Tyll et al., 2013; Werner & Noppeney, 2010). Although face-voice integration has been described in the auditory cortex (CL, CM, in awake and anesthetized monkeys; A1 only in awake monkeys) and the STS (Ghazanfar et al., 2008; Perrodin et al., 2015), and to a lesser extent in specific face-patches (Khandhadia et al., 2021), here, enhancement of the audio-visual response can only be seen in the blocked conditions involving visual scenes. The parsimonious interpretation of these observations is that face-vocalization binding was easier than scene-vocalization binding, thereby resulting in enhanced integrative processes, specifically in this latter condition, in agreement with the fact that neuronal multisensory integration is more pronounced for low saliency stimuli.

### Cortical and subcortical network for social audio-visual association based on meaning

In the present study, as indexed by the heart rate and hemodynamic brain signal modulation in the LS and the STS, the social visual stimulus used in each blocked condition sets the context and the subsequent distinctive processing of congruent versus incongruent auditory vocalization stimuli. We used a gPPI in order to identify the network contributing to setting this context. Both the LS and the STS are associated with a cortico-cortical network composed of the IPS, ACC (and vmPFC), DLPFC and OFC at the cortical level (and, to a lesser extent, the insula), and the amygdala and the hippocampus at the subcortical level.

The human brain has evolved a functional specialization for processing social information, such that the auditory cortex is involved in peri-lexical speech perception, visual areas in visual perception of speech, STS in faces and lips movement processing, the limbic system (amygdala, insula and ACC) in emotional processing and the IPS in spatially directed attention (Haxby et al., 2002; Haxby & Gobbini, 2011). A more recent review argues in favour of an interaction between attention and social processes to select information in a social environment (Capozzi & Ristic, 2018). This interaction is at play in three different levels of social processing: perception, interpretation and evaluation. First, attention acts at the perceptual level by facilitating relevant social information perception. Then a link with the emotional state of the individual and the social meaning of the cue is achieved, gating responses as a result of interpretation. Lastly, the valuation of the cue is estimated (Capozzi & Ristic, 2018). We hypothesize that the above-described network is homologous between macaques and humans, consisting of two interacting networks, one involved in the emotional processing of social stimuli, and one involved in their cognitive and attentional assessment.

We propose that the first homologous network involves the LS, the STS, the ACC (and vmPFC), the OFC, the amygdala, the hippocampus and the insula. This is in general agreement with the observation that species-specific vocalisations activate a network recruiting, in addition to the voice patches, visual areas such as V4, MT, STS areas TE and TEO, as well as areas from the limbic and paralimbic system, including the hippocampus, the amygdala and the ventromedial prefrontal cortex (vmPFC) (Gil-da-Costa et al., 2004). Brain stimulations applied to the auditory cortex directly activate vlPFC and indirectly the hippocampus (Rocchi et al., 2021). This is also in agreement with the finding that the observation of visual social interactions recruit vmPFC, vlPFC, ACC and OFC (Cléry et al., 2021; Roberts, 2006; Rudebeck et al., 2006; Rushworth et al., 2007; Sliwa & Freiwald, 2017).

We propose that the second homologous network involves the LS, the STS, the IPS and the DLPFC. DLPFC and area 46 are reciprocally connected with the caudal and rostral auditory cortex (Romanski et al., 1999). DLPFC is generally proposed to play a key role in attentional selection and memory processes (Courtney et al., 1997), and has been specifically associated with working memory during face processing (Rowe & Passingham, 2001). Spontaneous, non-trained responses to non-social (Guipponi et al., 2013; Schlack et al., 2005) and social (Joly, Pallier, et al., 2012; Ortiz-Rios et al., 2015; Poremba et al., 2003) auditory stimuli have been described in the IPS. In addition, the parieto-prefrontal network is classically associated with attentional selection (Buschman & Miller, 2007; Ibos et al., 2013) and forms with STS PITd area a larger attentional network (Sani et al., 2019). In humans, the IPS is described as a major node for the social and affective modulation by attention in naturalistic social visual information (see for review Frank & Sabatinelli, 2017), further interacting with the amygdala (for emotional processing), the hippocampus (for memory retrieval), and the OFC and PFC for top-down control over emotional processes. This is in agreement with our proposal of two homologous networks.

This homology opens the door to clinical research. Indeed, understanding these mechanisms is not only important from a comparative perspective with our own species, but may represent a fundamental contribution to issues concerning mental health. In particular, autistic individuals are often challenged by understanding social scenes, including the integration of auditory and visual information (Feldman et al., 2018; Stevenson, Siemann, Schneider, et al., 2014; Stevenson, Siemann, Woynaroski, et al., 2014a, 2014b). Such deficits may result from not only deficits in face and voice processing on their own, but the ability to integrate each modality in the service of predicting and understanding social interactions. This would implicate the two macaque networks we propose which give the possibility to test novel clinical approaches.

## Conclusion

**Our experiments demonstrate, using** indirect measures (heart rate and hemodynamic brain response), that macaque monkeys are able to associate social auditory and visual information based on their abstract meaning. This supports the idea that non-human primates display advanced social competences, amodally represented, that may have paved the way, evolutionary, for human social cognition. We further show that these processes recruit two functional networks that are, we propose, homologous to those observed in our own species.

## Contributions

Conceptualization, S.B.H. M.F; Stimuli preparation, M.H., M.F, Q.G, M.G; Data Acquisition, M.F. M.G.; Methodology, M.F., S.C., Q.G., and S.B.H; Investigation, M.F. and S.B.H.; Writing – Original Draft, M.F and S.B.H.; Writing – Review & Editing, S.B.H., M.F., M.H.; Funding Acquisition, S.B.H.; Supervision, S.B.H.

## Acknowledgements

S.B.H. were funded by the French National Research Agency (ANR)ANR-16-CE37-0009-01 grant and the LABEX CORTEX funding (ANR-11-LABX-0042) from the Université de Lyon, within the program Investissements d’Avenir (ANR-11-IDEX-0007) operated by the French National Research Agency (ANR). We thank Fidji Francioly and Laurence Boes for animal care, Julian Amengual and Justine Cléry for their rich scientific exchanges during data collection and analyses, Franck Lamberton and Danièle Ibarrola for their MRI methodological support and Holly Rayson for help on visual stimuli collection.

## Ethics declarations

The authors declare no competing interests. Animal experiments were authorized by the French Ministry for Higher Education and Research (project no. 2016120910476056 and 1588-2015090114042892) in accordance with the French transposition texts of Directive 2010/63/UE. This authorization was based on ethical evaluation by the French Committee on the Ethics of Experiments in Animals (C2EA) CELYNE registered at the national level as C2EA number 42.

## Material and methods

### Subjects and surgical procedures

Two male rhesus monkeys (*Macaca mulatta*) participated in the study (T, 15 years, 10kg and S, 12, 11kg). The animals were implanted with a Peek MRI-compatible headset covered by dental acrylic. The anaesthesia for the surgery was induced by Zoletil (Tiletamine-Zolazepam, Virbac, 5 mg/kg) and maintained by isoflurane (Belamont, 1–2%). Post-surgery analgesia was ensured thanks to Temgesic (buprenorphine, 0.3 mg/ml, 0.01 mg/kg). During recovery, proper analgesic and antibiotic coverage was provided. The surgical procedures conformed to European and National Institutes of Health Guidelines for the Care and Use of Laboratory Animals. The project was authorized by the French Ministry for Higher Education and Research (project no. 2016120910476056 and 1588-2015090114042892) in accordance with the French transposition texts of Directive 2010/63/UE. This authorization was based on ethical evaluation by the French Committee on the Ethics of Experiments in Animals (C2EA) CELYNE registered at the national level as C2EA number 42.

### Experimental setup

During the scanning sessions, monkeys sat in a sphinx position in a plastic monkey chair (Vanduffel et al., 2001) facing a translucent screen placed 60 cm from the eyes. Visual stimuli were retro-projected onto this translucent screen. Their head was restrained and the auditory stimuli were displayed by Sensimetrics MRI-compatible S14 insert earphones. The monkey chair was secured in the MRI with safety rubber stoppers to prevent any movement. Eye position (X, Y, right eye) was recorded thanks to a pupil-corneal reflection video-tracking system (EyeLink at 1000 Hz, SR-Research) interfaced with a program for stimulus delivery and experimental control (EventIDE®). Monkeys were rewarded for maintaining fixation into a 2×2° tolerance window around the fixation point.

### General run design

On each run, monkeys were required to fixate a central cross on the screen (Figure 1A). Runs followed a block design. Each run started with 10 s of fixation in the absence of sensory stimulation followed by three repetitions of a pseudo-randomized sequence containing six possible 16 s blocks: fixation (Fx), visual (Vi), auditory congruent (AC), auditory incongruent (AI), congruent audio-visual (VAC) and incongruent audio-visual (VAI). Each block (except the fixation block) consisted in an alternation of 500 ms stimuli (except for lip smacks, 1s dynamic stimuli succession) of the same semantic category (see Stimuli section below), in the visual, auditory or audio-visual modalities. Each block ended by 10 s of fixation in the absence of sensory stimulations.

### Face and social task design

Six audio-visual blocked conditions were presented to both monkeys, organized in runs as described above (Figure 1B). Six different blocked conditions were presented to both monkeys, organized in runs as described above (Figure 1B). Each task combined visual stimuli of identical social content with either semantically congruent or incongruent monkey vocalizations (Figure 1b). The face affiliative task (F+) combined lipsmaks with coos and aggressive calls. The face aggressive task (F-) combined aggressive faces with coos and aggressive calls. The first social affiliative task (S1+) combined grooming scenes with coos and aggressive calls. The second social affiliative task (S2+) combined grooming scenes with coos and screams. The social aggressive task (S1-) combined aggressive group or individual scenes with coos and aggressive calls. The social escape task (S2-) combined fleeing groups or individual scenes with coos and screams. Importantly, pairs of blocked conditions (F+ &. F-; S1+ & S1-; S2+ & S2-) shared the same auditory conditions, but opposite social visual content.

### Stimuli

Vocalizations were recorded from semi-free-ranging rhesus monkeys during naturally occurring situations by Marc Hauser. Detailed acoustic and functional analyses of this repertoire has been published elsewhere (e.g., Gouzoules et al., 1984; Hauser & Marler, 1993). Field recordings were then processed, restricting to selection of experimental stimuli to calls that were recorded from known individuals, in clearly identified situations, and that were free of competing noise from the environment. Exemplars from this stimulus set have already been used in several imaging studies(Belin et al., 2007; Cohen et al., 2007; Romanski, 2012; Romanski et al., 2005; Russ et al., 2008). All stimuli were normalized in intensity. The frequency ranges varied between the different types of stimuli as shown in Figure S4. For each of the three vocalization categories, we used 10 unique exemplars coming from matched male and female individuals, thus controlling for possible effects due to gender, social hierarchy or individual specificity. Coos are vocalisations typically produced during affiliative social interactions, including grooming, approach, coordinated movement, and feeding. Aggressive calls are typically used by a dominant animal toward a subordinate, often as a precursor to an actual physical attack. Screams are produced by subordinates who are either being chased or attacked, or as they are witnessing others in the same condition. Face (lipsmacks and aggressive facial expression) and social scene (group grooming, aggressive individual alone or in group / escaping individual or group) stimuli were extracted from videos collected by the Ben Hamed lab, as well as by Marc Hauser on Cayo Santiago, Puerto Rico. Images were normalized for average intensity and size. All stimuli were 4° x 4° in size. However, we decided to keep them in colour to get closer to natural stimuli even if it produced greater luminosity disparity between the different stimuli preventing us to use pupil diameter as a physiological marker. Only unambiguous facial expressions and social scenes were retained (Figure S4). A 10% blur was applied to all images, in the hope of triggering multisensory integration processes (but see result section). For each visual category, 10 stimuli were used.

### Scanning Procedures

The in-vivo MRI scans were performed on a 3T Magnetom Prisma system (Siemens Healthineers, Erlangen, Germany). For the anatomical MRI acquisitions, monkeys were first anesthetized with an intramuscular injection of ketamine (10 mg\kg). Then, the subjects were intubated and maintained under 1-2% of isoflurane. During the scan, animals were placed in a sphinx position in a Kopf MRI-compatible stereotaxic frame (Kopf Instruments, Tujunga, CA). Two L11 coils were placed on each side of the skull and a L7 coil was placed on the top of it. T1-weighted anatomical images were acquired for each subject using a magnetization-prepared rapid gradient-echo (MPRAGE) pulse sequence. Spatial resolution was set to 0.5 mm, with TR= 3000 ms, TE=3.62 ms, Inversion Time (TI)=1100 ms, flip angle=8°, bandwidth=250 Hz/pixel, 144 slices. T2-weighted anatomical images were acquired per monkey, using a Sampling Perfection with Application optimized Contrasts using different flip angle Evolution (SPACE) pulse sequence. Spatial resolution was set to 0.5 mm, with TR= 3000 ms, TE= 366.0 ms, flip angle=120°, bandwidth=710 Hz/pixel, 144 slices. Functional MRI acquisitions were as follows. Before each scanning session, a contrast agent, composed of monocrystalline iron oxide nanoparticles, Molday ION™, was injected into the animal’s saphenous vein (9-11 mg/kg) to increase the signal to noise ratio (Leite et al., 2002; Vanduffel et al., 2001). We acquired gradient-echoechoplanar images covering the whole brain (TR=2000 ms; TE=18 ms; 37 sagittal slices; resolution: 1.25×1.25×1.38 mm anisotropic voxels) using an eight-channel phased-array receive coil; and a loop radial transmit-only surface coil (MRI Coil Laboratory, Laboratory for Neuro- and Psychophysiology, Katholieke Universiteit Leuven, Leuven, Belgium, see Kolster et al., 2014). The coils were placed so as to maximise the signal on the temporal lobe.

### Data description

In total, 76 runs were collected in 12 sessions for monkey T and 65 runs in 9 sessions for monkey S. Based on the monkey’s fixation quality during each run (85% within the eye fixation tolerance window) we selected 60 runs from monkey T and 59 runs for monkey S in total, i.e. 10 runs per task, except for one task of monkey S.

### Data analysis

Data were pre-processed and analysed using AFNI (Cox, 1996), FSL (Jenkinson et al., 2012; Smith et al., 2013), SPM software (version SPM12, Wellcome Department of Cognitive Neurology, London, UK, https://www.fil.ion.ucl.ac.uk/spm/software/), JIP analysis toolkit (http://www.nitrc.org/projects/jip) and Workbench (https://www.humanconnectome.org/software/get-connectome-workbench). The T1-weighted and T2-weighted anatomical images were processed according to the HCP pipeline (Autio et al., 2020; Glasser et al., 2013) and were normalized into the MY19 Atlas (Donahue et al., 2016). Functional volumes were corrected for head motion and slice time and skull-stripped. They were then linearly realigned on the T2-weighted anatomical image with flirt from FSL, the image distortions were corrected using nonlinear warping with JIP. A spatial smoothing was applied with a 3-mm FWHM Gaussian Kernel.

Fixed effect individual analyses were performed for each monkey, with a level of significance set at p<0.05 corrected for multiple comparisons (FWE, t-scores 4.6) and p<0.001 (uncorrected level, t-scores 3.09). Head motion and eye movements were included as covariate of no interest. Because of the contrast agent injection, a specific MION hemodynamic response function (HRF) (Vanduffel et al., 2001) was used instead of the BOLD HRF provided by SPM. The main effects were computed over both monkeys. In most analyses, face blocked conditions and social blocked conditions were independently pooled.

ROI analyses were performed as follows. ROIs were determined from the auditory congruent contrast (AC vs Fx) of face blocked conditions with the exception of two ROIs of the right lateral sulcus (LS4 and LS6) that were defined from the same contrast of social blocked conditions. ROIs were defined as 1.5 mm diameter spheres centred around the local peaks of activation. In total, 8 ROIs were selected in the right STS, 6 from the left STS, 4 in the left LS and 6 in the right LS. Figure S1 shows the peak activations defining each selected ROI; so as to confirm the location of the peak activation on either of the inferior LS bank, the superior STS bank or the inferior STS bank. For each ROI, the activity profiles were extracted with the Marsbar SPM toolbox (marsbar.sourceforge.net) and the mean percent of signal change (+/- standard error of the mean across runs) was calculated for each condition relative to the fixation baseline. %PSC were compared using Wilcoxon non-parametric paired tests.

Generalized Form of Context-Dependent Psychophysiological Interactions was performed as follows (gPPI, http://www.nitrc.org/projects/gppi), using the CONN toolbox (www.nitrc.org/projects/conn, RRID:SCR_009550), an open-source Matlab/SPM-based cross-platform software. Specifically, we were interested in identifying the network activated throughout the runs and that could account for the dependence of auditory perception on the visual context of the task. The gPPI analysis was performed independently on each task, over the averaged LS and STS ROIs time series respectively, for each hemisphere. We report, in supplementary table 1 the cortical and subcortical regions the time series of which showed a significant temporal correlation with the seeds (p=0.005 uncorrected). Are considered for discussion only the cortical regions the time series of which correlate, with either the LS or STS time series, in at least three of the six blocked conditions, for each of the left and right hemispheres (criterion 1), or in at least three of the six blocked conditions, in at least one hemisphere and in at least eight of the blocked conditions across both hemispheres (criterion 2, more stringent).

### Behaviour and Heart rate

During each run of acquisition, videos of the faces of monkeys S and T were recorded in order to track heart rate variations (HRV) as a function of blocked conditions and blocks (Froesel et al., 2020). We focus on heart rate variations between auditory congruent and incongruent stimuli. For each task, we extracted HRV during AC and AI blocs. As changes in cardiac rhythm are slow, analyses were performed over the second half (8s of each block). This has been done for each run of each task, grouping both monkeys. Because the data were not normally distributed (Kolmogorov-Smirnov Test of Normality), we carried out Friedman tests and non-parametric post hoc tests.

## Supplementary Material

**Figure S1:**
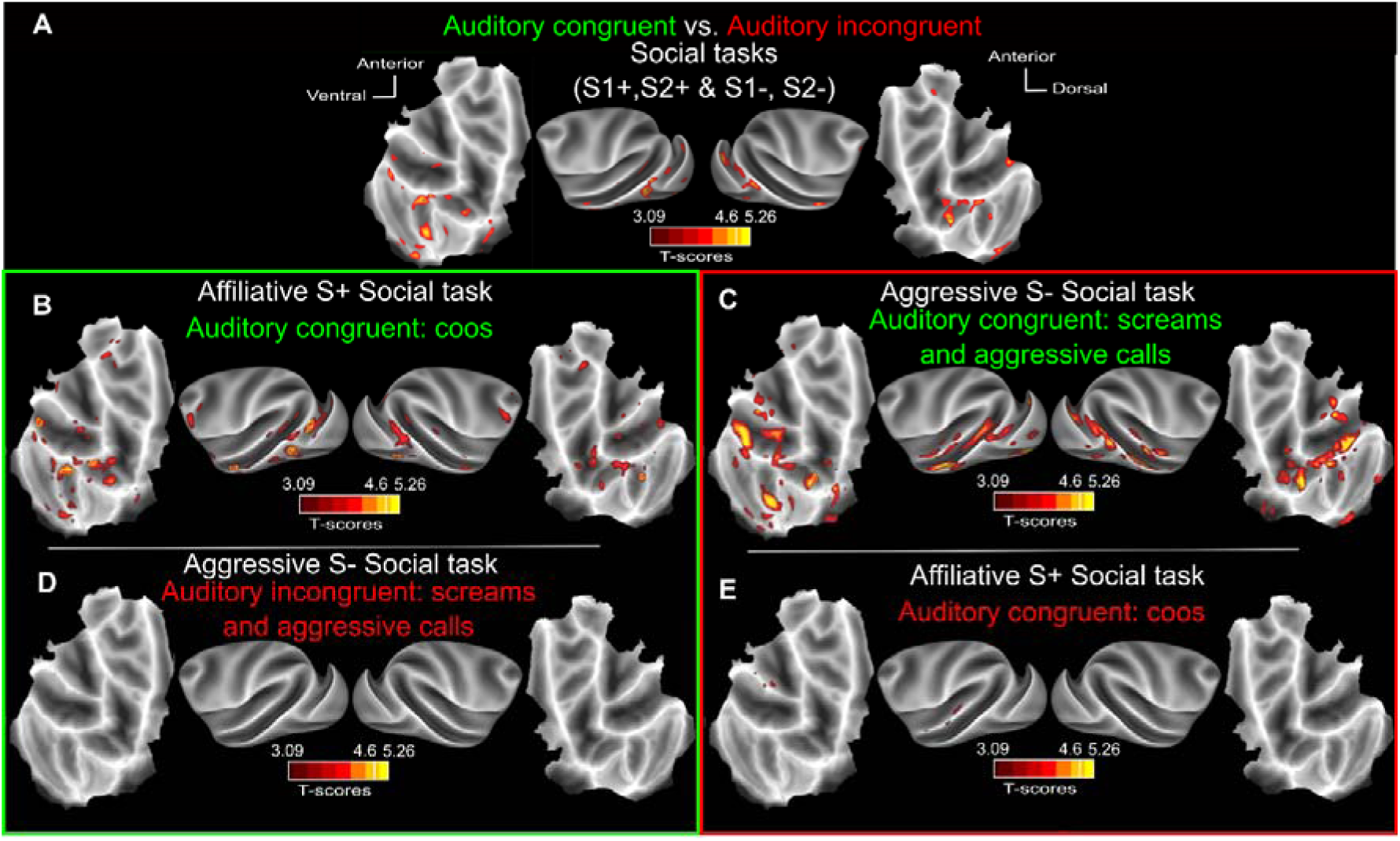
**Auditory activations depend on semantic congruence with visual context in social blocked conditions.** A) Whole-brain activation maps of the S1+, S2+ (social affiliative 1 & 2), S1- (social aggressive) and S2- (social escape) runs, for the auditory congruent vs auditory incongruent (relative to the visual context) contrast. B) Whole-brain activation map for the S+ (social affiliative, S1+&S2+) auditory congruent (coos, dark green, AC vs. Fx) and auditory incongruent (aggressive calls and screams, dark red, AI vs. Fx) conditions. C) Whole-brain activation map for the S- (social negative, S1-&S2-) auditory congruent (aggressive calls and screams, dark green, AC vs. Fx) and auditory incongruent (coos, dark red, AI vs. Fx) conditions. Darker shades of red indicate level of significance at p<0.001 uncorrected, t-score 3.09. Lighter shades of yellow and brown outlines indicate level of significance at p<0.05 FWE, t-score 4.6.

**Figure S2:**
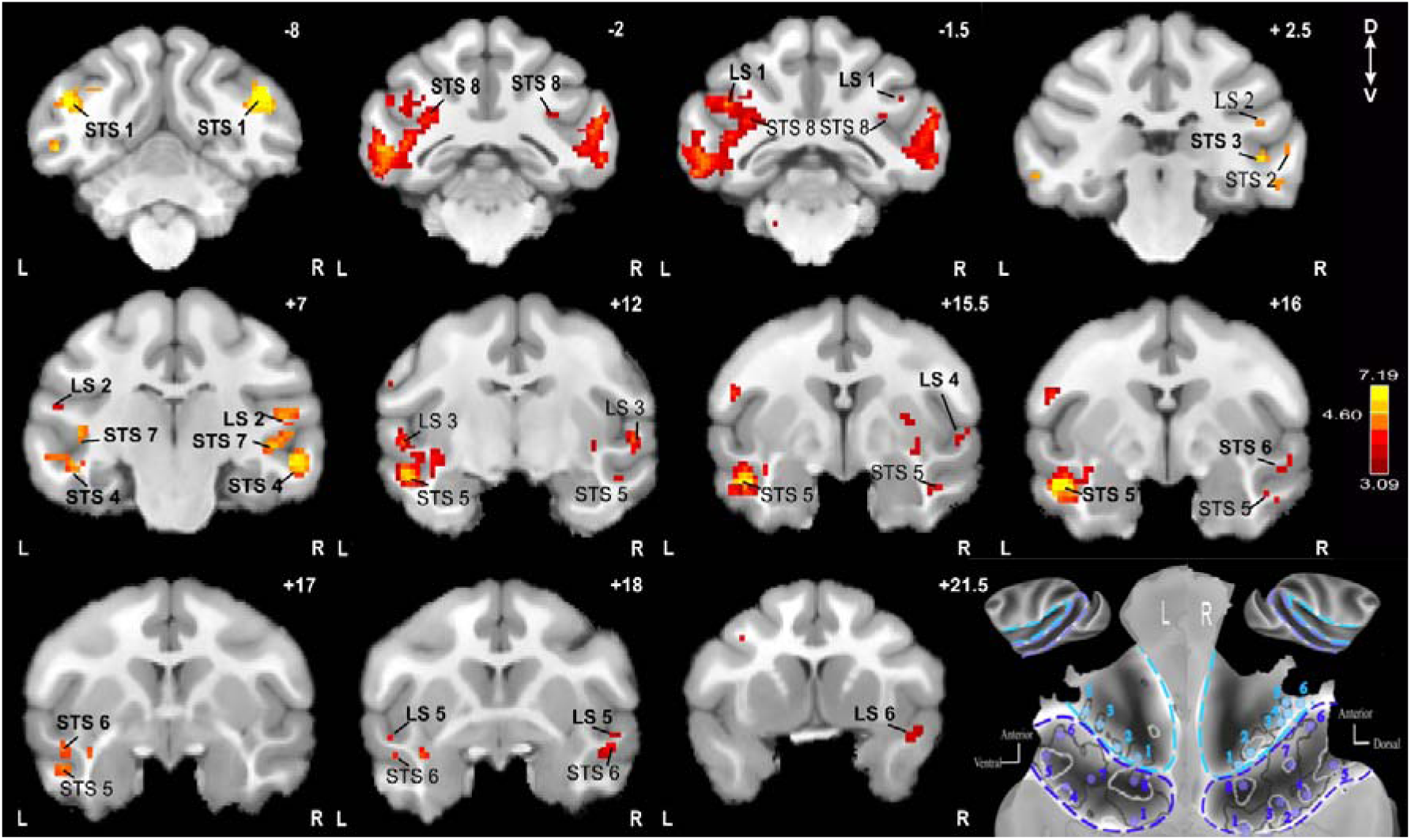
**Peak activations that were used to define the ROIs of interest, on coronal slices, based on the AC vs. Fx contrast presented in (Figure 2 and Figure 4).** Antero-posterior level relative to the intra-aural line indicated at the top right corner of each slice, in millimeters. Bold fonts refer to the location peak. Normal fonts refer to cortical activation extending beyond the peak. Activation thresholds were varied on some sections in order to clearly show the existence of local activation maxima. Activation color-scale was however kept constant across all slices. Down-left panel: ROIs locations on flatmaps (lateral sulcus; light blue; superior temporal sulcus: dark blue), same conventions as in Figure 5.

**Figure S3:**
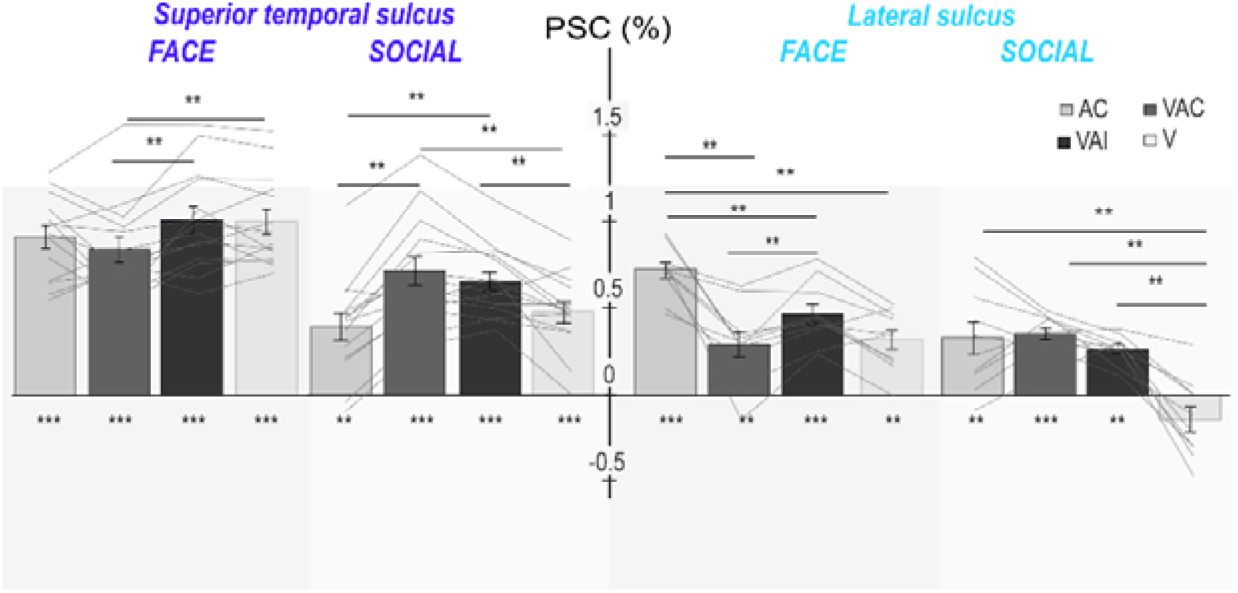
**Percentage of signal change (%PSC) across all lateral sulcus (light blue) and superior temporal sulci (dark blue) ROIs of both hemispheres, comparing unimodal and multimodal congruent and incongruent conditions.** Statistical differences relative to fixation are between conditions are indicates as follows: **, p<0.01 (Wilcoxon non-parametric test).

**Figure S4:**
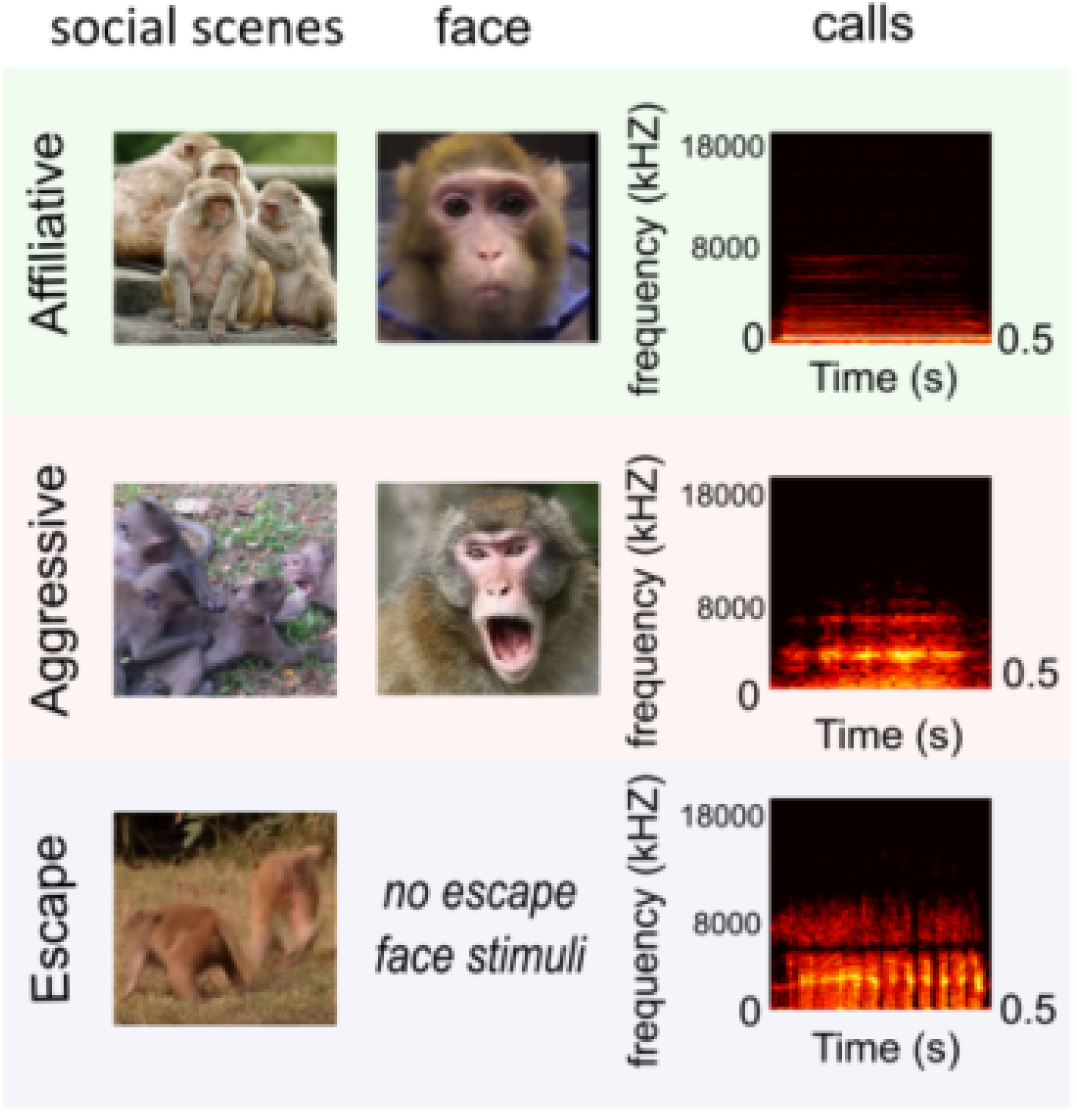
**Example of visual and auditory stimuli.** At the right are shown examples of visual stimuli used for social and face blocked conditions. On the left, congruent calls spectrograms are associated to the visual stimuli are shown. The affiliative call is a coos, the aggressive congruent auditory stimulus is an aggressive call and the escape call is a scream. Stimuli were not strictly normalized in terms of in low visual and auditory feature properties, thus making their social meaning the dominant cue across the different stimuli of a given category.

**Table S1:** **Whole brain gPPI functional connectivity analysis for the left and right lateral sulcus and the left and right superior temporal sulcus ROIs.** Highlighted are the cortical regions showing at least one voxel with significant correlation with seed regions (p<0.005 uncorrected).

| Left LS | IPS | Insula | dIPFC (area 46) | STS | ACC | OFC | hippocampus | amygdala | dorsal pulvinar |
| --- | --- | --- | --- | --- | --- | --- | --- | --- | --- |
| F+ | + | + | + | + | - | - | - | + | - |
| F- | + | + | + | + | + | + | - | + | - |
| S1- | + | - | + | + | + | + | + | + | - |
| S2- | + | - | + | + | + | + | - | - | - |
| S1+ | + | + | + | + | + | + | + | + | + |
| S2+ | - | + | + | + | + | + | + | - | + |

| Right LS | IPS | Insula | dIPFC (area 46) | STS | ACC | OFC | hippocampus | amygdala | dorsal pulvinar |
| --- | --- | --- | --- | --- | --- | --- | --- | --- | --- |
| F+ | + | + | + | + | - | + | - | + | - |
| F- | + | - | + | + | - | + | - | + | - |
| S1- | + | - | + | + | + | + | + | + | + |
| S2- | + | - | + | + | + | - | + | + | + |
| S1+ | - | - | + | + | + | + | + | - | - |
| S2+ | + | + | + | + | + | + | + | + | + |

| Left STS | IPS | Insula | dIPFC (area 46) | LS | ACC | OFC | hippocampus | amygdala | dorsal pulvinar |
| --- | --- | --- | --- | --- | --- | --- | --- | --- | --- |
| F+ | + | - | + | + | - | + | + | + | - |
| F- | - | + | + | + | + | + | + | - | + |
| S1- | + | - | - | + | + | + | + | - | + |
| S2- | + | + | - | + | + | + | - | + | + |
| S1+ | + | + | + | + | + | + | - | - | - |
| S2+ | + | - | + | + | - | - | + | + | - |

| Right STS | IPS | Insula | dIPFC (area 46) | LS | ACC | OFC | hippocampus | amygdala | dorsal pulvinar |
| --- | --- | --- | --- | --- | --- | --- | --- | --- | --- |
| F+ | + | + | - | + | + | + | - | + | + |
| F- | + | + | + | - | + | + | + | + | - |
| S1- | + | - | + | + | + | - | - | + | - |
| S2- | + | + | - | - | + | + | - | - | - |
| S1+ | + | - | - | + | + | + | + | + | - |
| S2+ | + | + | + | + | + | + | + | + | + |

## Notes

### Competing Interest Statement

The authors have declared no competing interest.

